# Immune Response of Transplanted Kidney Tissues Assembled from Organoid Building Blocks

**DOI:** 10.1101/2023.08.26.551822

**Authors:** Thiago J. Borges, Yoshikazu Ganchiku, Jeffrey O. Aceves, Ronald van Gaal, Sebastien G. M. Uzel, Jonathan E. Rubins, Kenichi Kobayashi, Ken Hiratsuka, Murat Tekguc, Ivy A. Rosales, Guilherme T. Ribas, Karina Lima, Rodrigo B. Gassen, Ryuji Morizane, Jennifer A. Lewis, Leonardo V. Riella

**Affiliations:** Center for Transplantation Sciences, Massachusetts General Hospital, Harvard Medical School, Boston, MA; Paulson School of Engineering and Applied Sciences, Harvard University, Cambridge, MA; Wyss Institute for Biologically Inspired Engineering, Harvard University, Boston, MA; Nephrology Division, Department of Medicine, Massachusetts General Hospital, Harvard Medical School, Boston, MA; Harvard Stem Cell Institute (HSCI), Cambridge, MA; Department of Pathology, Massachusetts General Hospital and Harvard Medical School, Boston, MA

## Abstract

The increasing scarcity of organs and the significant morbidity linked to dialysis requires the development of engineered kidney tissues from human-induced pluripotent stem cells. To accomplish this, integrative approaches that synergize scalable kidney organoid differentiation, tissue biomanufacturing, and comprehensive assessment of their immune response and host integration are essential. Here, we create engineered human kidney tissues composed of kidney organoid building blocks (OBBs) and transplant them into mice reconstituted with allogeneic human immune cells. We assess their host vascular integration, *in vivo* maturation, and their ability to trigger human immune responses. Tissue-infiltrating human immune cells are composed of effector T cells and innate cells. This immune infiltration leads to kidney tissue injury characterized by reduced microvasculature, enhanced kidney cell apoptosis, and a unique inflammatory gene signature comparable to kidney organ transplant rejection in humans. Upon treatment with the immunosuppressive agent Rapamycin, the induced immune response is greatly suppressed. Our model serves as a translational platform to study engineered kidney tissue immunogenicity and develop novel therapeutic targets for kidney rejection.

## Introduction

The kidneys are vital organs that maintain homeostasis by controlling fluid and electrolyte balance, removing waste products, and regulating blood pressure. In the U.S. alone, roughly 25 million people have chronic kidney disease (CKD) (NIH, 2023; United States Renal Data System). Many of these patients will progress to end-stage renal disease (ESRD), for which kidney transplantation is the best treatment. However, due to the lack of transplantable organs, patients typically require dialysis. Although dialysis is capable of helping with fluid and electrolyte balance (Teitelbaum, 2021), it is associated with substantial morbidity and mortality primarily due to cardiovascular disease (Aufhauser et al., 2018; Teitelbaum, 2021). The five-year survival rate of people on hemodialysis is less than 50% — worse than some types of cancer (United States Renal Data System). By contrast, kidney transplantation increases patient survival, reduces costs, and improves their quality of life (Schnuelle et al., 1998; Vollmer et al., 1983). In the U.S. alone, more than 650,000 people are on dialysis and ∼95,000 people await kidney transplants with over 3,000 new patients added monthly to the wait list (http://optn.transplant.hrsa.gov). Given the growing shortage of organs and the substantial morbidity associated with dialysis, there is a need to develop alternative sources of kidney tissue that can restore normal kidney function.

We postulate that biomanufacturing functional kidney tissues, and, ultimately, whole organs may offer an important solution to this growing problem. Towards this goal, engineered human kidney tissues must be scalably produced and successfully integrated with the host upon transplantation. Recently, differentiation protocols have been developed to generate “mini-kidneys” from human-induced pluripotent stem cells (hiPSCs) (Morizane and Bonventre, 2017; Morizane et al., 2015; Taguchi et al., 2014; Takasato et al., 2015). These organoids exhibit remarkable tissue microarchitectures with high cellular density akin to their *in vivo* counterparts. Integrative approaches that combine scalable organoid differentiation with tissue biomanufacturing as well as the evaluation of immune responses and host integration are needed to begin to close the gap from organoid building blocks (OBBs) to therapeutic organs (Wolf et al., 2022).

Here, we created engineered human kidney tissue assembled from kidney organoid building blocks (OBBs) and assessed their *in vivo* vascular integration, maturation, and elicited immune responses in a humanized mouse model. Kidney organoids were differentiated from the human induced pluripotent stem cells, suspended in a fibrinogen solution, compacted into a dense tissue matrix, and then patterned into thin tissue discs (3 mm in diameter, 1 mm in thickness, with a total volume of ∼10 mm^3^). Their final composition is akin to the living tissue matrices used for a bioprinting method known as sacrificial writing into functional tissue (SWIFT) (Skylar-Scott et al., 2019; Wolf et al., 2022). iPS cells were HLA-typed and kidney tissues were transplanted into the fat pad of NSG mice reconstituted with allogeneic human immune cells. Analysis of the infiltrating human immune cells within the kidney tissues revealed the presence of activated CD4^+^ and CD8^+^ T cells as well as innate cells, such as monocytes and natural killer (NK) cells. This correlated with a distinctive inflammatory gene signature within the transplanted kidney tissues. Subsequently, we observed transplant rejection features characterized by an increase in kidney apoptotic cells and reduced tissue vascularization and maturation markers. However, treatment with the immunosuppressive agent, Rapamycin, effectively inhibited the human immune response and prevented kidney tissue injury. Our integrated platform enables the investigation of HLA-incompatible immune responses to human-engineered human organoid-based tissues, as well as the efficacy and nephrotoxicity of new therapeutical agents.

## Results

### Engineering kidney tissues from organoid building blocks

Human kidney tissues are generated by creating kidney organoids differentiated from hiPSCs by following our previously published protocol of directed differentiation (Morizane and Bonventre, 2017; Morizane et al., 2015), with modifications for stirred bioreactors or spinner flasks that enable 3D differentiation (**Fig 1A**). This protocol involves differentiation into metanephric mesenchyme, as confirmed by SIX2 immunostaining on day 8 of differentiation. The resulting kidney organoids exhibit glomerular and tubular features under brightfield imaging (**Fig 1B**), staining positive for podocyte (podocalyxin-like protein 1, PODXL), proximal tubule (lotus tetragonolobus lectin, LTL), distal tubule (cadherin 1, CDH1), and endothelial (platelet endothelial cell adhesion molecule, PECAM1/CD31) markers (**Fig 1C-D**). The kidney organoids are then suspended in a fibrinogen-based (10 mg/ml) extracellular matrix solution and compacted via centrifugation to form a densely cellular tissue matrix, which is cast into a mold containing gelatin and thrombin. The mold is pre-patterned using a 3D printed stamp to produce a periodic array of 18 cylindrical features (diameter = 3 mm and height = 1mm, **Fig. 1E**). The gelatin and thrombin mold is initially cast at 37°C and features are produced by adding the stamp, cooling to 4°C and removing the stamp, leaving the desired cavities. Upon casting the organoid-fibrinogen solution into these cylindrical molds, the thrombin diffuses from the surrounding extracellular gelatin matrix to promote its rapid polymerization to fibrin, which initially binds the organoids together prior to their fusion (**Fig 1F**). The kidney tissue discs are cultured for 7 days to promote organoid fusion prior to implantation. This process leads to a pronounced reduction in tissue disc diameter (**Fig 1G-J**) as well as a diminishing presence of fibrin over this period (**Fig 1K-N**). Importantly, these kidney tissue discs continue to exhibit robust expression of glomerular, proximal tubular, and early stages of microvessel formation, as reflected by CD31 expression (**Fig 1O-R**).

**Figure 1.**
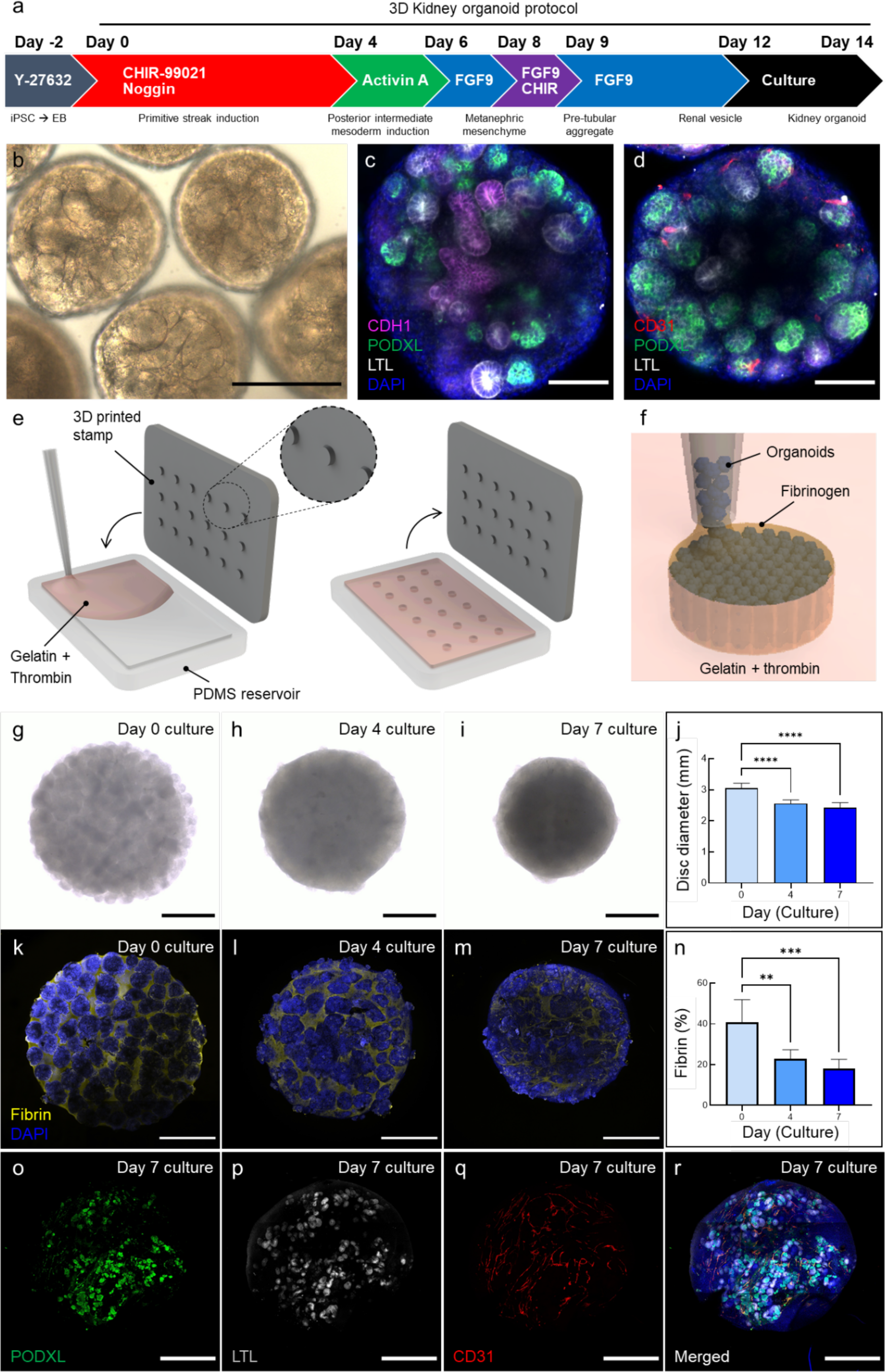
Generation and characterization of human kidney tissues assembled from organoid building blocks. **(A)** 3D differentiation protocol used to generate kidney organoids from hiPSCs. **(B)** Brightfield image of kidney organoids differentiated and cultured in spinner flasks (day 14), scalebar = 200 µm. **(C-D)** Confocal images of kidney organoids differentiated and cultured in spinner flasks (day 14), confirming expression of podocyte (PODXL, green), proximal tubule (LTL, gray), distal tubule (CDH1, magenta), and endothelial (CD31, red) cell populations, scale bars = 100 µm. **(E-F)** Schematic illustrations of kidney tissue disc fabrication method, in which a 3D printed stamp is molded into a PDMS reservoir that contains a gelatin and thrombin mixture to pattern cylindrical features of diameter = 3 mm and height = 1 mm. Kidney tissue discs are produced by filling these molded features with a densely cellular assembly of kidney organoids suspended in a fibrinogen solution, which rapidly polymerizes into fibrin via diffusion of thrombin from the surrounding gelatin matrix. **(G-I)** Optical images of the kidney tissue discs cultured at days 0, 4, and 7 on an orbital shaker, scale bars = 1 mm. **(J)** Plot of kidney tissue disc diameter as a function of culture time, which reflects organoid fusion into a contiguous tissue, 2-way ANOVA, *** = p<0.001. **(K-M)** Confocal images showing the reduction in fibrin (yellow) content as a function of culture time (days 0, 4, and 7) for these kidney tissue discs, scale bars = 1 mm. **(N)** Corresponding plot of fibrin (yellow) content as a function of culture time for these tissues, which reflects the measured area fraction of fibrin determined by image analysis, 2-way ANOVA, ** = p<0.01, *** = p<0.001. After 7 days of static culture and tissue fusion, these kidney organoid-based tissues still contained **(O)** podocyte (PODXL, green), **(P)** proximal tubule (LTL, gray), and **(Q)** endothelial (CD31, red) cells that self-organize into **(r)** glomerular, tubular, and microvascular architectures, respectively, scale bars = 1 mm.

### Immune response to the transplanted engineered kidney tissues

To investigate the immune responses triggered by the kidney organoid-based tissue, we subcutaneously transplanted kidney organoid tissue discs (∼10 mm^3^) into the fat pad of the immunocompromised NSG mice. On the following day, the transplanted NSG mice were reconstituted with allogeneic human immune cells (peripheral blood mononuclear cells, PBMCs) isolated from a healthy volunteer or PBS 1x (controls). We then evaluated the immune response in the kidney discs and spleens over time (**Fig 2A**). Prior to transplantation, the iPSCs and immune cells were HLA typed, and four HLA mismatches on A, B and DR loci (A1, B2 and DR1) were identified at the antigen level (**Suppl. Table 1**). To improve the precision of the alloimmune risk assessment, we also characterized the donor-recipient HLA mismatch at the molecular level (Sypek and Hughes, 2021). Our data revealed 41 eplet mismatches in HLA class I and 23 in class II.

**Figure 2.**
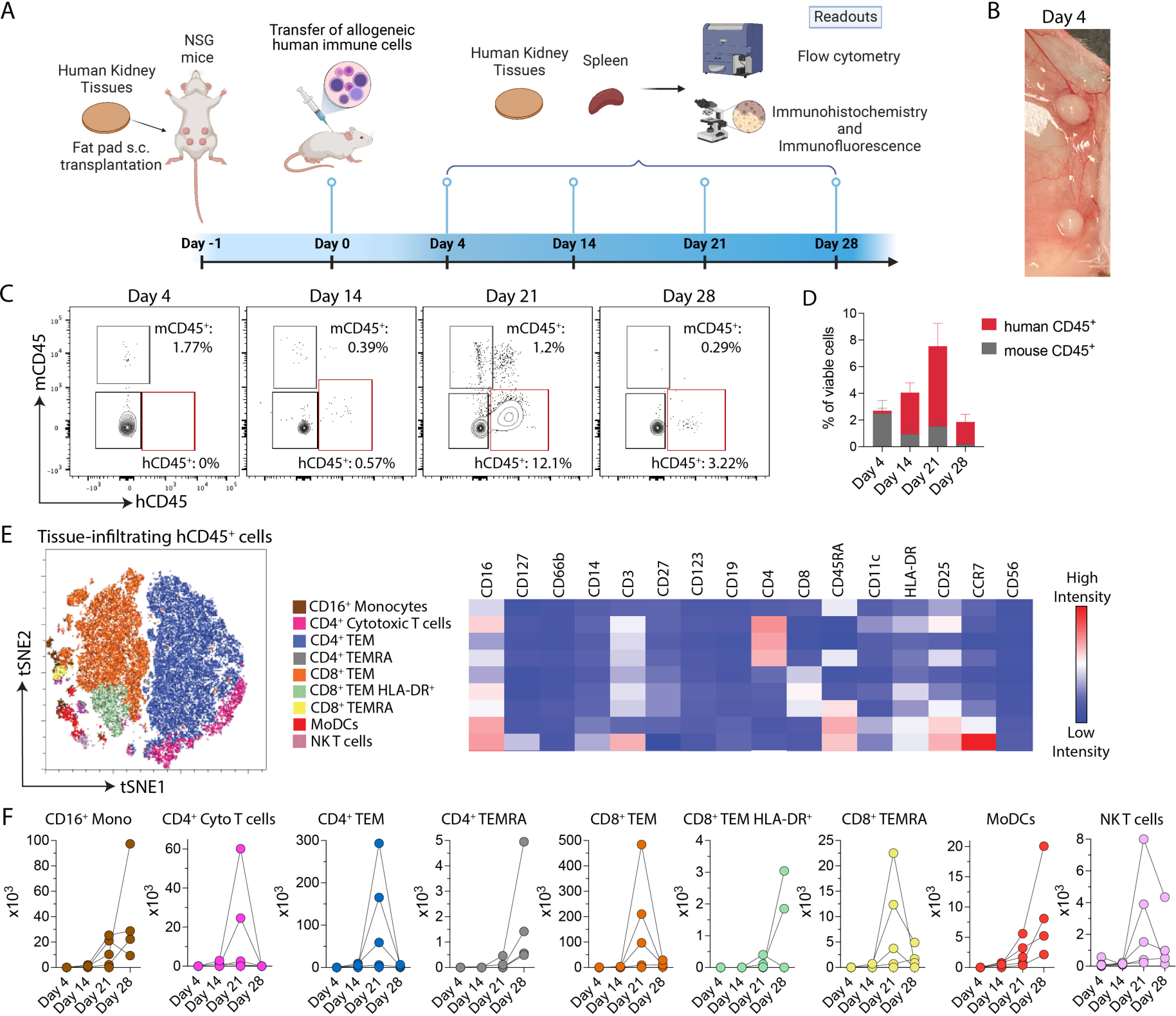
Immune response to transplanted kidney tissues. (**A**) Kidney tissues were subcutaneously (s.c.) transplanted into the fat pad of NSG mice reconstituted with or without allogeneic human immune cells (the source is freshly isolated PBMCs). The kidney tissues were recovered and analyzed by histology and flow cytometry on days 4, 14, 21 and 28 after allogeneic human immune cells were transferred. (**B**) Image of the kidney tissues harvested on day 4. (**C**) Representative contour plots (gated on viable singlets) and (**D**) percentages of mouse and human kidney tissue-infiltrating CD45^+^ leukocytes over time. (**E**) The viSNE plot and heatmap represent the expression of immune markers in nine immune cell meta clusters identified by FlowSOM. (**F**) Absolute numbers of tissue-infiltrating human CD16^+^ monocytes, CD4^+^ cytotoxic T cells, CD4^+^ T effector memory (TEM), CD4^+^ T effector memory CD45RA+ (TEMRA), CD8^+^ TEM, CD8^+^HLA-DR^+^TEM, monocyte-derived dendritic cells (MoDCs), and natural killer (NK) T cells on days 4, 14, 21 and 28 after the transfer of allogeneic human immune cells. Data are representative of two pooled experiments (n = 3-5 tissues/group/time point).

At the tissue scale, we observed host integration and vascularization of transplanted kidney tissues harvested on days 4, 14, 21 and 28 after the transfer of allogeneic human immune cells (**Fig 2B**, **Suppl. Fig 1A**). There was no difference between the gross weights of discs harvested from NSG mice with or without an allogeneic human immune system (**Suppl. Fig 1B**). We next evaluated the infiltration of human and mouse CD45^+^ leukocytes over time. Our data demonstrated that human immune cell infiltration peaked within these kidney tissues at day 21 post-PBMC injection (**Fig 2C-D**). T cells are the major drivers of kidney transplant rejection (Montgomery et al., 2018). To characterize the subsets of the human immune cells infiltrating the transplanted tissues, we performed an unsupervised hierarchical clustering by employing in-depth immune phenotyping with an 18-marker flow cytometry panel. We identified 9 tissue-infiltrating human immune cell populations, with the majority of cells composed of T cells (**Fig 2E**). Within the T cell populations, we found that CD4^+^ cytotoxic (CD16^+^) T cells, CD4^+^ and CD8^+^ T effector memory (TEM), and CD8^+^ T effector memory that express CD45RA (TEMRA) peaked on day 21 after the transfer of allogeneic human immune cells (**Fig 2F**). From the innate compartment, we identified CD16^+^ monocytes and monocyte-derived dendritic cells (MoDCs) populations that increased over time, peaking at day 28 (**Fig 2F**). Similar to the effector T cells, the absolute number of natural killer (NK) T cells increased on day 21 (**Fig 2F**). The increased infiltration of CD8^+^ and CD4^+^ mononuclear cells over time was confirmed by immunohistochemistry (**Suppl. Fig. 2A-B**). Importantly, all observed immune subsets were associated with an effector/activated phenotype. Overall, our findings indicate that the transplantation of human kidney tissues assembled from organoid building blocks in the presence of allogeneic human immune cells is capable of generating effector innate and adaptive immune responses against the allo-kidney tissue.

### In vivo expression of kidney markers by transplanted tissues

The *in vivo* viability of kidney organoid building blocks is an important attribute for their use in creating engineered tissues, and, ultimately, transplantable organs (Ibi and Nishinakamura, 2023). Hence, we evaluated the nephron epithelia of our transplanted tissues in the absence or presence of allogeneic human immune cells. Histologically, increasing interstitial inflammation is observed on days 21 and 28 after the transfer of allogeneic human immune cells (**Fig 3A**). Immunohistochemical studies using podocalyxin (PODXL, podocytes) and E-cadherin (CDH1, distal tubule) demonstrated that our transplanted kidney tissue discs exhibited glomerular (PODXL^+^, **Fig 3B**) and tubular CDH1^+^, **Fig 3C**) structures that were maintained over time in the absence of allogeneic human immune cells. However, in the presence of allogeneic human immune cells, these structures were degraded by day 28 (**Fig 3B, C)**. Our data clearly show that transplanted kidney tissues remain intact *in vivo* after implantation in NSG mice, while the introduction of allogeneic human immune cells leads to injury of glomerular and tubular structures.

**Figure 3.**
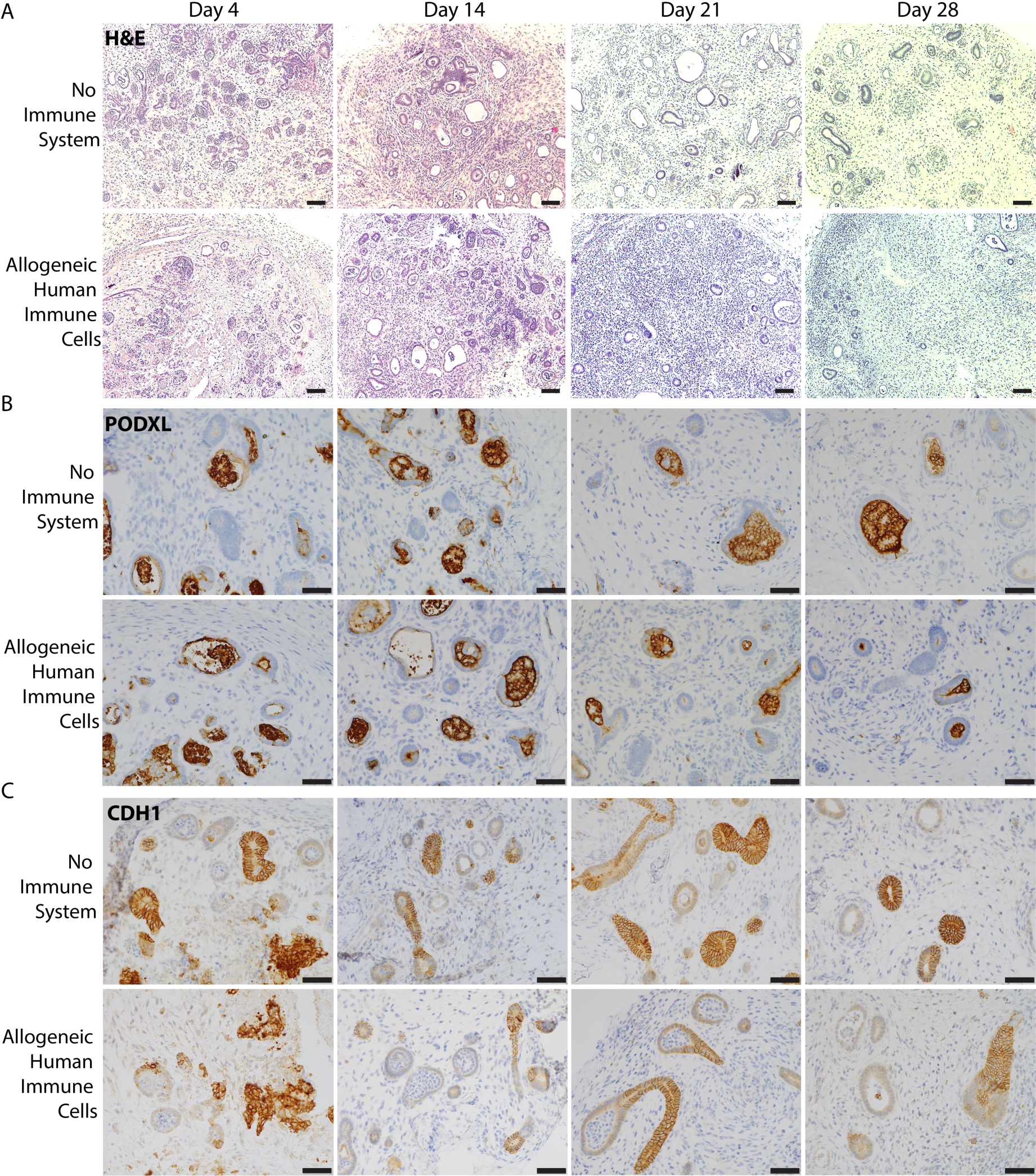
Transplanted tissues express markers of different kidney cell subsets. Kidney tissues were transplanted in NSG mice reconstituted with or without allogeneic human immune cells. (**A**) Representative disc H&E staining, and immunohistochemistry analysis of kidney tissues. Immunohistochemical analyses of (**B**) PODXL (glomeruli) and (**C**) CDH1 (distal tubules) expression on days 4, 14, 21 and 28 after the transfer of allogeneic human immune cells. Data are representative of three independent experiments. Scale bars = 50 μm.

### Allogeneic human immune cells induce engineered kidney tissue rejection

During transplantation, the recipient’s immune system recognizes donor HLA molecules as foreign and initiates a rejection response (Montgomery et al., 2018). To evaluate kidney tissue immunogenicity following *in vivo* transplantation, we stained the harvested tissues for HLA class I (HLA-A, B and C) and II (HLA-DR). We found that the kidney tissues expressed HLA I in the absence of allogeneic human immune cells over time (**Fig 4A**). By contrast, HLA II was not detected in the group without immune cells (**Fig 4B**). In the presence of allogeneic human immune cells, HLA class I immunostaining increased (**Fig 4A**), including expression by interstitial infiltrating mononuclear cells. Furthermore, in NSG mice reconstituted with allogeneic human immune cells, harvested kidney tissues exhibited increased interstitial HLA-DR^+^ immune cells, reaching a peak on day 21 (**Fig 4B**). These data confirm our flow cytometry findings regarding the infiltration of both antigen-presenting cells and HLA-DR^+^ T cells within these transplanted kidney tissues (**Fig 2D and E**). While HLA class II molecules are primarily expressed by antigen-presenting cells, kidney tubules can upregulate HLA class II in the setting of injury (Zhou et al., 2021). Indeed, we observed tubular expression of HLA-DR on days 21 and 28 after the transfer of allogeneic human immune cells (**Fig 4B, arrow**). Using transcriptomics, we evaluated the discs on day 21 by NanoString and confirmed an enhanced presence of HLA class I (HLA-A, B, C), and HLA class II (HLA-DQA1, DRB1 and DRB5) transcripts in tissues transplanted in mice reconstituted with allogeneic human immune cells compared to controls (**Fig 4C**).

**Figure 4.**
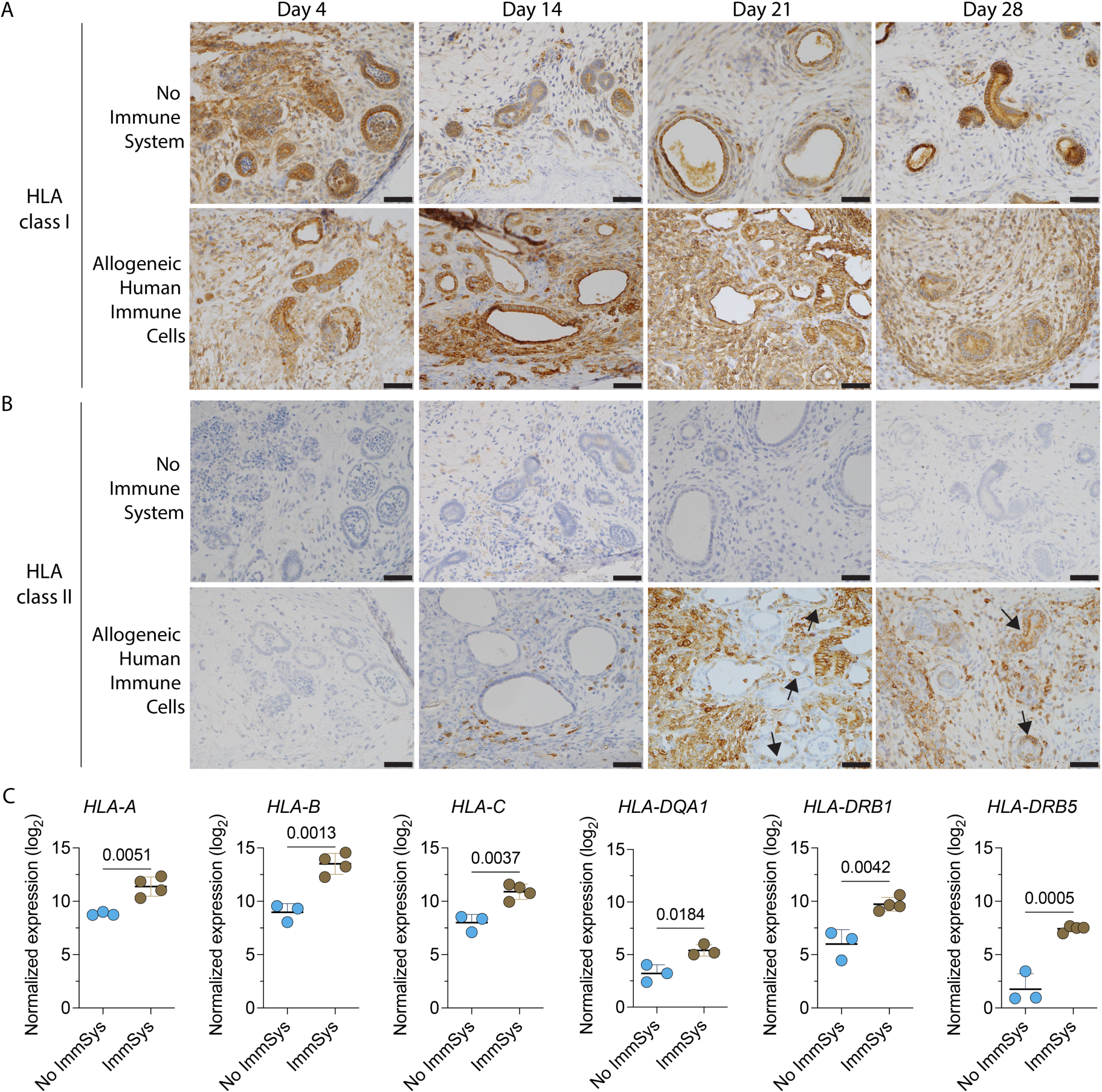
Kidney organoid-based tissues express HLA class I and II *in vivo*. Kidney tissues are transplanted into NSG mice reconstituted with or without allogeneic human immune cells, harvested and then analyzed on days 4, 14, 21 and 28. Immunostaining of kidney tissues (**A**) HLA class I (HLA-A, B and C), and (**B**) HLA class II (HLA-DR) expression over time. Data are representative of two independent experiments. Arrows indicate tubules expressing HLA-DR. Scale bars = 50 μm. (**C**) Log_2_ normalized gene expression of disc *HLA-A*, *HLA-B*, *HLA-C*, *HLA-DQA1*, *HLA-DRB1*, and *HLA-DRB5* transcripts from NSG mice with no immune system (No ImmSyst, n = 3) and reconstituted with allogeneic human immune cells (ImmSyst, n = 4) by NanoString analysis. Statistical analyses are done by unpaired t-tests.

To assess host integration and vascularization in the absence or presence of allogeneic human immune cells, we stained and quantified the endothelial marker CD31 in these transplanted kidney tissues. We found that the number of CD31^+^ vessels increased over time in the absence of immune cells by immunohistochemistry (**Fig 5A**) and immunofluorescence **(Fig 5B)**. By contrast, the CD31^+^ vessels decreased in tissues exposed to allogeneic human immune cells (**Fig 5A**), suggesting enhanced injury to the vascular endothelium. Confocal imaging confirmed the decrease of CD31^+^ cells and maturation markers in kidney tissues harvested from NSG mice reconstituted with allogeneic human immune cells (**Fig 5B, Suppl Fig 3A**). Next, we used the TUNEL assay to detect DNA-strand breaks, which are characteristic of apoptotic cells (Kepp et al., 2011). Apoptosis of cells within the transplanted kidney tissues increased markedly in NSG mice reconstituted with allogeneic human immune cells (**Suppl. Fig 3B**). We further confirmed these findings by staining the harvested tissues with Annexin-V to assess apoptosis in the non-immune cell population by flow cytometry. The percentage of Annexin-V^+^ cells increased over time and was significantly higher in the discs harvested from NSG mice reconstituted with allogeneic human immune cells compared to controls (**Fig 5C, D**). Taken together, our results reveal that bioengineered kidney tissues express HLA molecules upon *in vivo* implantation, enhancing their immunogenicity. These data also show that our humanized mouse model is capable of triggering transplant rejection of these engineered kidney tissues.

**Figure 5.**
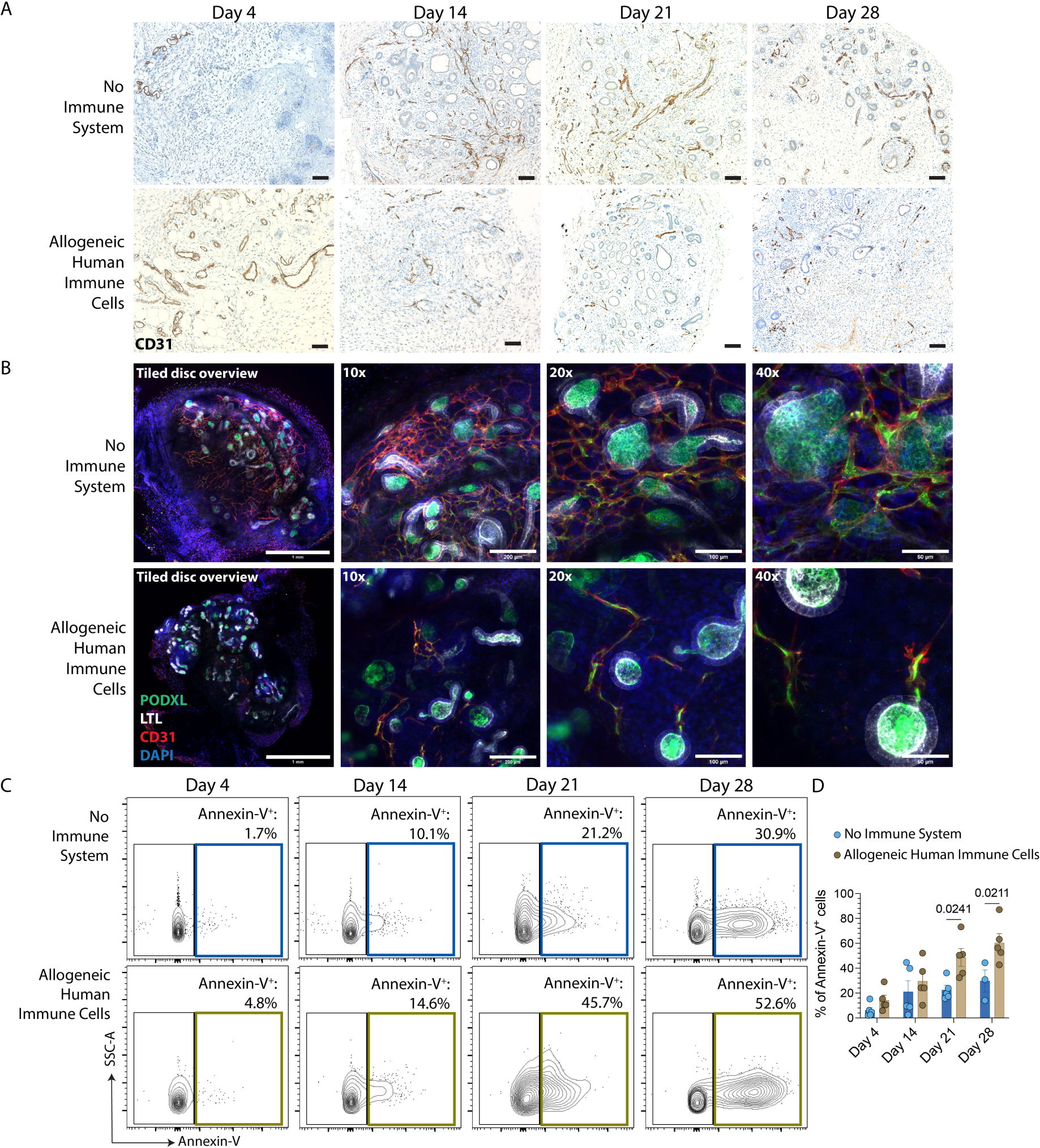
Transplanted kidney organoid-based tissues are rejected by allogeneic human immune cells *in vivo*. Kidney tissues are transplanted into NSG mice reconstituted with or without allogeneic human immune cells, harvested and then analyzed on days 4, 14, 21 and 28. (**A**) Immunostaining of disc CD31 (endothelial cells) expression over time. Scale bars = 100 μm. Data are representative of three independent experiments (**B**) Confocal images confirming the expression of PODXL (glomeruli, green), LTL (proximal tubules, gray), CD31 (endothelium) and DAPI (nuclei, blue) by tissues harvested on day 21. Scale bars = 1 mm (tiled disc overview), 200 µm (10x), 100 µm (20x), and 50 µm (40x). Data are representative of two independent experiments. (**C)** Representative contour plots and (**D**) quantification of Annexin-V^+^ (apoptotic) cells by flow cytometry on kidney tissues harvested over time. Statistical analyses are done by two-way ANOVA followed by Šídák’s multiple comparison tests (n = 3-5 tissues/group/time point).

### In vivo kidney tissue rejection is inhibited by an immunosuppressive drug

The control of effector rejection responses with immunosuppressive drugs is crucial for the successful clinical translation of engineered tissues, and, ultimately whole organs. Indeed, the development of models that mimic human kidney transplant rejection provides useful platforms for assessing novel therapeutic targets. Hence, we used our model to evaluate the effects of the immunosuppressive agent Rapamycin (Rapa, sirolimus) in kidney tissue injury induced by allogeneic human immune cells. Inhibitors of the mTOR pathway, like Rapa, are widely used as a class of maintenance immunosuppressive drugs after organ transplantation (Bestard et al., 2007; Knechtle et al., 2003), and they have the benefit of not only inhibiting effector T cells but also of promoting regulatory T cells (Battaglia et al., 2005). We transplanted kidney tissues in NSG mice reconstituted with allogeneic human immune cells and administered Rapa daily (**Fig 6A**). Transplanted discs were analyzed by flow cytometry, histology and NanoString on day 21, at the peak of their immune response. Host integration and vascularization were observed in all cases (**Suppl. Fig 4A**), with their weights decreased in tissues harvested from NSG mice treated with Rapa (**Suppl. Fig 4B**). Human and CD45^+^ leukocytes were also significantly decreased in the kidney tissues (**Fig 6B**) and spleens (**Fig 6C**) of NSG mice reconstituted with allogeneic human immune cells and treated with Rapa. These mice had smaller spleens with reduced weight (**Suppl. Fig 4C-D**), suggesting an inhibition of the alloimmune response. The immune characterization of the human leukocytes infiltrating the transplanted tissues demonstrated that Rapa treatment significantly decreased tissue-infiltrating monocytes (**Fig 6D, E**), total and activated CD8^+^ T cells (**Fig 6D, F**), MoDCs (**Fig 6D, G**), and NK T cells (**Fig 6D, H**). Interestingly, this treatment also increased the proportion of Tregs in the tissues (**Fig 6D, I**). Overall, our findings demonstrate that Rapa treatment can control innate and adaptive effector immune responses and proportionally increases Tregs *in vivo*.

**Figure 6.**
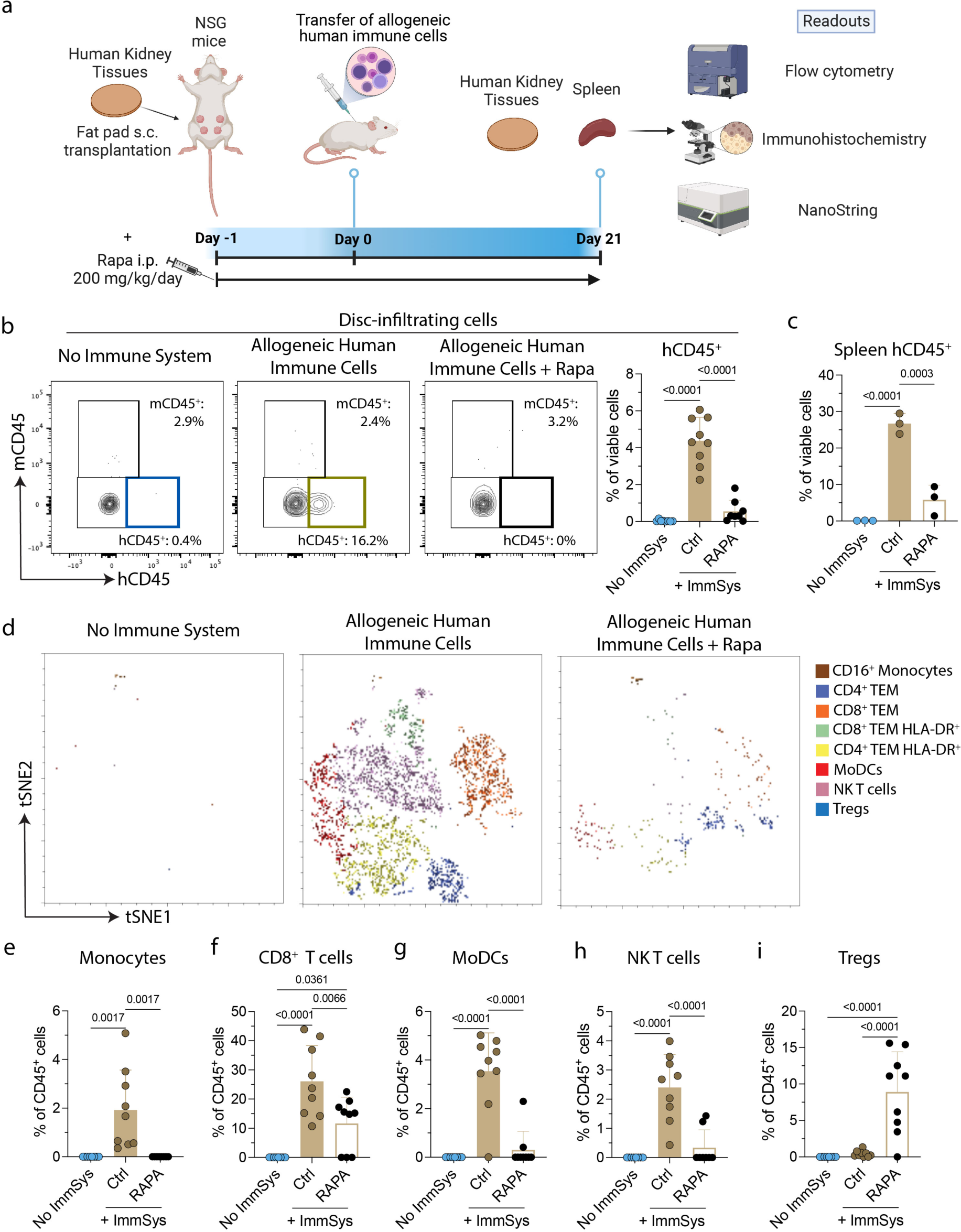
Rapamycin controls kidney tissue-effector cell immune response *in vivo*. (**A**) Kidney tissues were subcutaneously (s.c.) transplanted into the fat pad of NSG mice reconstituted with or without allogeneic human immune cells. One cohort of NSG mice was intraperitoneally (i.p.) treated with 200 mg/kg/day of Rapamycin (Rapa). The kidney tissues were recovered and then analyzed by histology, flow cytometry and NanoString analysis on day 21. (**B, C**) Representative contour plots (gated on viable singlets) and percentages of the (**B**) kidney tissue-infiltrating cells and (**C**) splenic human CD45^+^ leukocytes. Data are representative of two independent experiments. Statistical analyses are done by one-way ANOVA with Tukey post-test (n = 7-9 tissues/group, and 3 spleens/group). (**D**) viSNE plots represent the eight immune cell meta-clusters identified by FlowSOM in the three groups. Percentages of disc-infiltrating (**E**) human monocytes, (**F**) total CD8^+^ T cells, (**G**) monocyte-derived dendritic cells (MoDCs), (**H**) natural killer (NK) T cells, and (**I**) regulatory T cells (Tregs) at day 21 after the transfer of allogeneic human immune cells. Data are representative of two independent experiments. Statistics are performed by one-way ANOVA with Tukey’s multiple comparisons test (n = 7-9 tissues/group).

Next, we investigated whether Rapa treatment could interfere with host vascularization and inhibit tissue cell death induced by the allogenic human immune cells. Rapa significantly improved the numbers of CD31^+^ vessels in the kidney tissues (**Fig 7A, B**), reducing vascular endothelium injury. Apoptosis of kidney cells in these tissues was also significantly decreased in Rapa-treated mice as confirmed by TUNEL (**Fig 7A**) and Annexin-V (**Fig 7C**) stains. The observed changes in the phenotypes of infiltrating immune cells and suppression of kidney injury mimic the immunomodulatory effects of Rapa in the clinical setting. Our data suggest that the use of immunosuppressive drugs, like Rapamycin, can prevent the rejection of the transplanted kidney tissues assembled from organoid building blocks.

**Figure 7.**
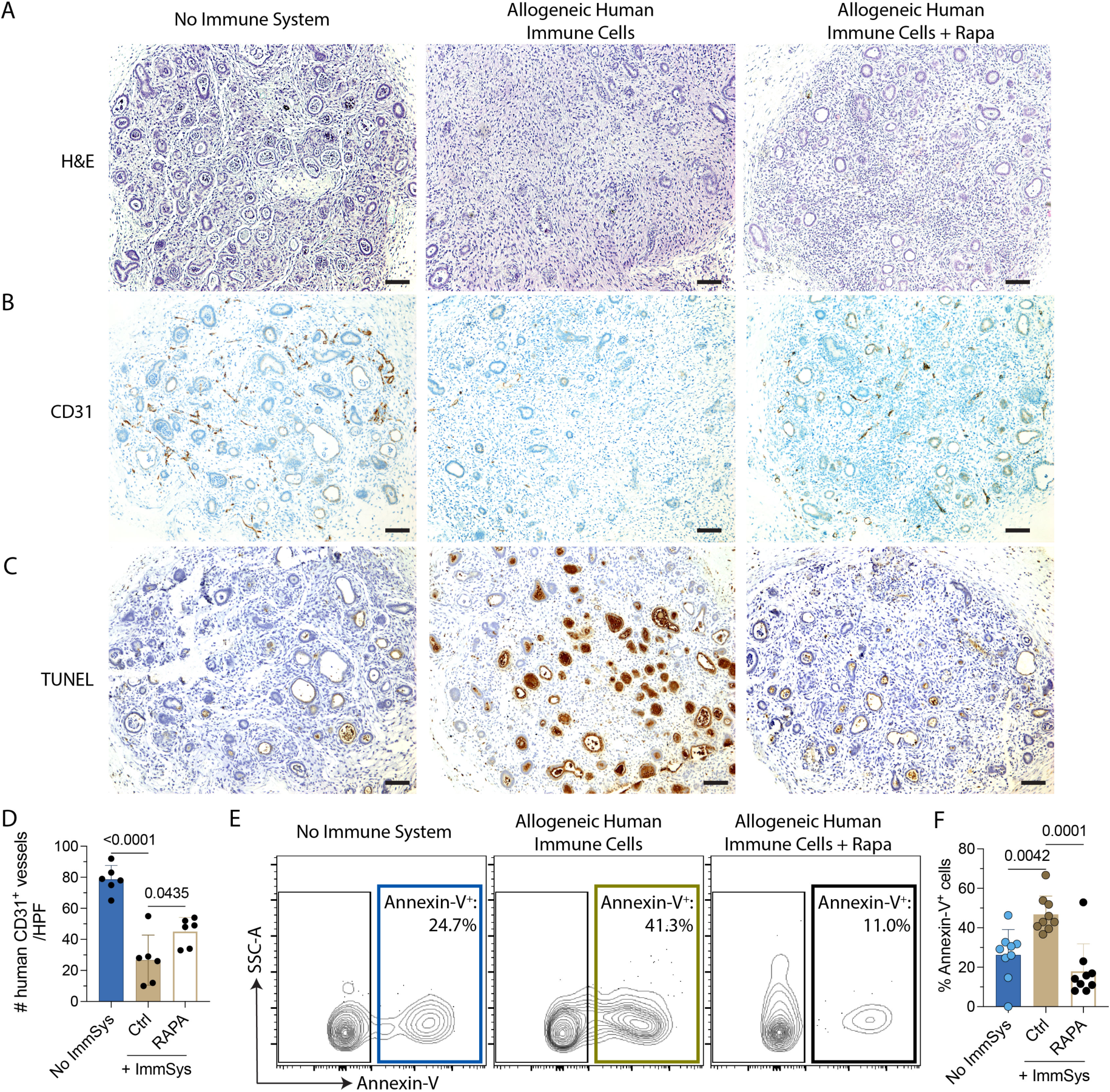
Rapamycin controls immune rejection of transplanted kidney organoid-based tissues. NSG mice were kidney tissue discs-implanted as displayed in Figure 6A. Tissues are harvested and analyzed on day 21 after the transfer of allogeneic human immune cells. (**A**) Representative H&E staining, (**B**) immunostaining of CD31, and (**C**) TUNEL assay in the three experimental groups. Data are representative of three independent experiments. Scale bars = 100 μm. (**D**) Quantification of disc CD31 staining. The number of CD31+ vessels was quantified from two representative 400x fields from each transplanted disc (n = 3). Statistic by one-way ANOVA with Tukey post-test (n = 6 tissues/group). (**E**) Representative contour plots and (**F**) quantification of Annexin-V^+^ (apoptotic) cells by flow cytometry on kidney discs harvested on day 21 after the transfer of allogeneic human immune cells. Statistic by one-way ANOVA with Tukey’s multiple comparisons test (n = 9 tissues/group).

### Characterizing the kidney tissue microenvironment during rejection

To determine the molecular characteristics associated with kidney rejection, we compared the gene expression profiles of kidney tissues harvested from NSG mice with no immune system, or reconstituted with allogeneic human immune cells treated or not with Rapa. Among 770 genes analyzed using the NanoString technology, 166 genes were differentially expressed during rejection (log2 fold change > 1.5; adjusted p-value < 0.05). The top 50 differentially expressed genes (DEGs) are shown in **Fig 8A**. The complete list of DEGs is displayed in **Supp. Table 2**. Of the 166 DEGs, 148 were upregulated in the allogeneic human immune cells group and included molecules related to allograft rejection, antigen processing and presentation, and TNF signaling pathway (**Fig 8B**).

**Figure 8.**
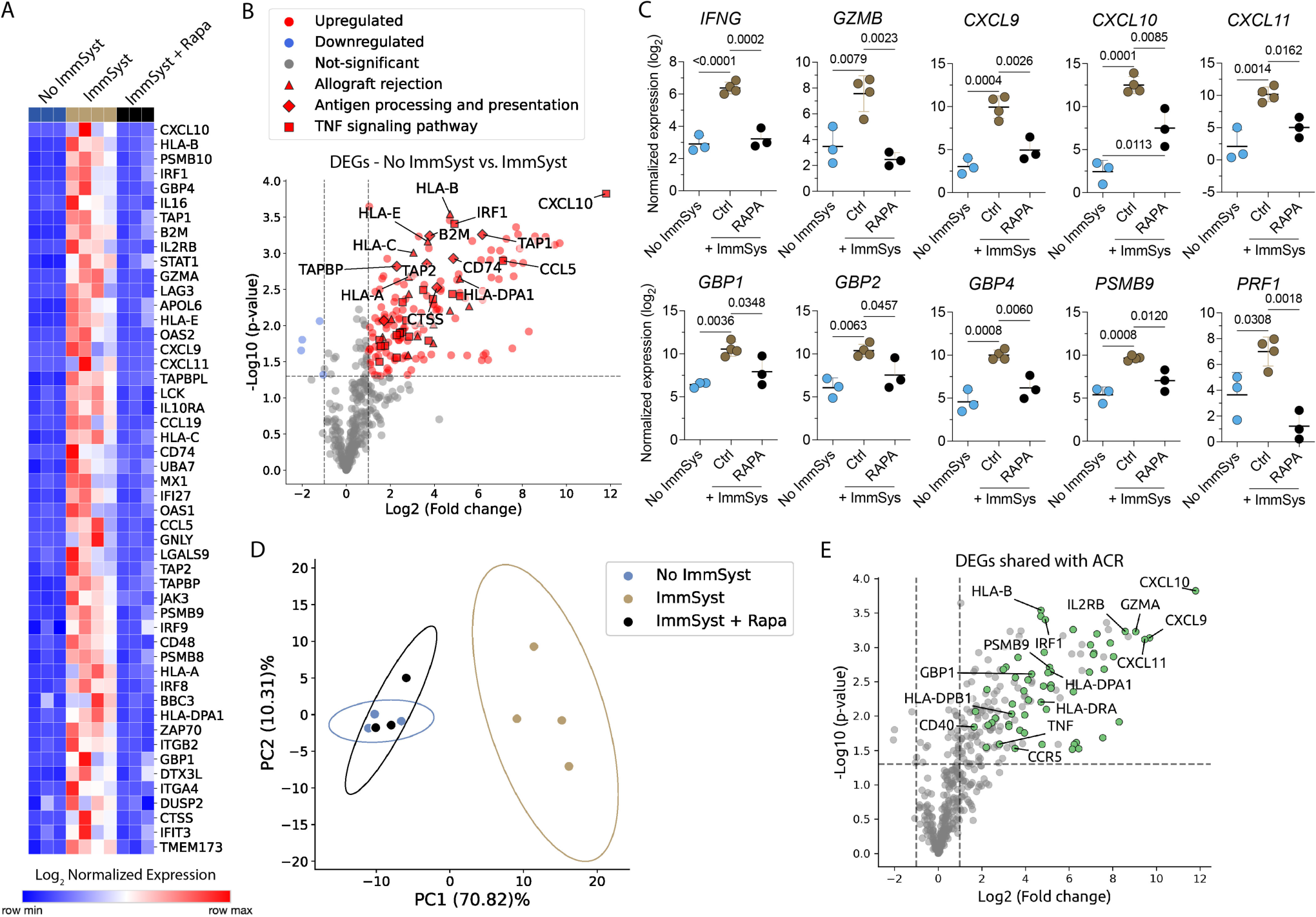
Unique rejection-related gene expression signature of transplanted kidney organoid-based tissues harvested from NSG mice reconstituted with allogeneic human immune cells. Kidney tissues were transplanted into NSG mice reconstituted with (ImmSyst) or without (No ImmSyst) allogeneic human immune cells. One cohort of NSG mice was treated with 200 mg/kg/day of Rapamycin (ImmSyst + Rapa). The kidney tissues were recovered and then analyzed by Nanostring on day 21. (**A**) Heatmap of the top 50 differentially expressed genes (DEGs) in tissues from NSG mice reconstituted with allogeneic human immune cells (ImmSyst, n = 4) and those treated with Rapa (ImmSyst + Rapa, n = 3) compared with NSG mice with no immune system (No ImmSyst, n = 3). Data expressed as log_2_ normalized expression. (**B**) Volcano plot showing DEGs in the group ImmSyst in relation to the No ImmSyst group. Each dot represents an individual gene, and each shape represents a KEGG pathway. Log_2_ fold change is represented on the x-axis, and the y-axis displays log_10_ of each gene’s p-value. (**C**) Log_2_ normalized gene expression of IFN-γ-related transcripts in the No ImmSyst, ImmSyst and ImmSyst + Rapa groups. Statistics used one-way ANOVA with Tukey’s multiple comparison test. (**D**) Unsupervised principal-component analysis of the top 166 DEGs, clustering the samples from NSG mice reconstituted with allogeneic human immune cells (ImmSyst) system compared with No ImmSyst and ImmSyst + Rapa events, which clustered together. (**E**) Volcano plots showing DEGs shared with human kidney transplant acute cellular rejection. Log_2_ fold change is represented on the x-axis, and the y-axis displays log_10_ of each gene’s p-value.

In kidney transplant recipients, cellular rejection is associated with a gene signature related to IFN-γ responses (Mengel et al., 2020; Reeve et al., 2009). We analyzed whether the presence of allogeneic human immune cells would induce the expression of IFN-γ−related genes in the kidney tissues and how Rapa could modulate their expression. We found an increase in the expression of kidney tissues *IFNG*, *GZMB*, *CXCL9*, *CXCL10*, *CXCL11*, *GBP1*, *GBP2*, *GBP4*, *PSMB9*, and *PRF1* after the transfer of allogeneic human immune cells (**Fig 8C**). The expression of these transcripts was significantly reduced by the treatment with Rapa (**Fig 8C**). A principal component analysis (PCA) conducted using all 166 DEGs revealed distinct clustering. The gene signature of the kidney tissues harvested from mice with allogeneic human immune cells differed from those without an immune system. Notably, transplanted kidney tissues from NSG mice without an immune system and those reconstituted with allogeneic human immune cells and treated with Rapa clustered together (**Fig 8D**), confirming its ability to reduce effector immune responses. Finally, we compared the gene expression profile of the rejecting kidney tissues with molecular signatures of acute cellular rejection in human kidney transplants. As previously reported by our group (Win et al., 2021), mRNA transcripts measured in kidney biopsy specimens during acute rejection were used. We found that the transcriptional profile of our kidney organoid-based tissues resembled the gene signature observed in human kidney transplant rejection (**Fig 8E**), especially an IFN-γ−related signature.

## Discussion

The engineering and transplantation of functional human kidney tissue could provide a potential alternative for treating patients with end-stage renal disease. We developed a fabrication method of kidney tissue discs comprising multiple kidney organoids, which were then transplanted subcutaneously into the fat pad of immunocompromised NSG mice. These tissues express PODXL (podocytes), CDH1 (distal tubule), LTL (proximal tubule), and CD31 (endothelial cells) *in vitro* and are able to engraft and survive *in vivo*. The presence of allogeneic human immune cells had harmful effects on the transplanted kidney tissues, including reducing their inherent vascularization (decreased expression of the CD31 endothelial marker) and enhancing cell death. A decrease in nephron epithelial markers, e.g., PODXL and CDH1, was also observed when transplanted kidney tissues were exposed to allogeneic human immune cells. Importantly, the rejection process exhibited similar molecular characteristics to those observed during kidney transplant rejection in humans. Importantly, rejection was suppressed upon administering the immunosuppressant, rapamycin. Our model serves as a platform for studying the immunogenicity of engineered transplanted tissues and new immunosuppressant drugs under development for kidney transplant recipients.

Kidney replacement therapy consists of life-supporting interventions to restore kidney function in patients with ESRD. While kidney transplantation is the most effective kidney replacement therapy, an insufficient supply of transplantable donor kidneys requires the majority of patients with renal failure to undergo dialysis, which mechanically filters waste products and excess fluids from the blood. However, dialysis is associated with severe morbidity and mortality, in particular, cardiovascular disease (Aufhauser et al., 2018; Vollmer et al., 1983). Numerous strategies are being explored for the development of transplantable kidneys, including xenotransplantation, decellularization and recellularization of scaffolds, kidney organoid-based therapies, and interspecies blastocyst complementation (Ibi and Nishinakamura, 2023). Kidney organoids derived from patient-specific human induced pluripotent stem cells are already serving as valuable tools for studying kidney development and diseases. Our study provides a foundational step towards using kidney organoids as building blocks for biomanufacturing transplantable kidney tissues, which may restore partial or complete function.

By understanding the role of HLA mismatches that serve as the main antigens triggering alloimmune responses during transplantation, we can create engineered kidney tissues that are more robust to rejection. While HLA molecules have lower expression in kidney organoids *in vitro*, we observed an increased expression of both class I and II upon transplantation and in the presence of allogeneic human immune cells. Moreover, transcriptional analyses of the kidney tissues in NSG mice reconstituted with allogeneic human immune cells revealed a similar molecular signature as observed in biopsies of kidney transplant recipients with acute cellular rejection with dominant IFN-γ-related transcripts (Mengel et al., 2020; Reeve et al., 2009). Furthermore, evidence of target kidney tissue damage was evident, confirming the immune-mediated injury. We found that rapamycin inhibits the effector alloimmune response and migitates kidney tissue injury. It should be noted that our model only mimics acute cellular rejection, which is the dominant mode of rejection during the first year post-transplantation, since PBMCs do not fully recapitulate all components of the human immune system, such as the engraftment of B cells and antibody production. Looking ahead, we plan to implant engineered kidney tissues with embedded vascular and drainage networks and assess their *in vivo* function.

In summary, our ability to scalably create kidney organoid-based tissues has advanced our understanding of the immunological challenge of transplanting HLA-incompatible tissues. Our observations underscore the importance of applying therapeutic approaches to minimize this alloimmune response via the use of immunosuppressive drugs or autologous or hypoimmunogenic stem cells for organoid and tissue generation.

## Methods

### hPSC culture and kidney organoid differentiation

BJFF human-induced PSCs (provided by Prof. Sanjay Jain at Washington University) are maintained in either StemFit02/04 or mTeSR plus stem cell media and used up to passage 50. For maintenance, BJFF iPSCs were passaged upon reaching ∼50 - 70% confluency. BJFFs are seeded into new plates/flasks coated with 1% Matrigel in DMEM/F12 medium, as previously described (Morizane and Bonventre, 2017). BJFFs are lifted from the original plate/flask by removing the stem cell medium, washing 1X with PBS without Ca^2+^ and Mg^2+^, incubating with Accutace or ReLeSR (STEMCELL Technologies) for 5-10 min at 37°C, and then using stem cell media to collect the lifted iPS cells. The BJFFs are then seeded into the new, pre-coated, plates/flasks in StemFit02/04 or mTeSR plus medium. Medium is changed no later than every 48-72h.

Before starting kidney organoid differentiation, BJFFs are grown to 80% confluency and lifted into a single cell suspension by removing the stem cell medium, washing 1X with PBS without Ca^2+^ and Mg^2+^, and incubating with Accutace or ReLeSR (STEMCELL Technologies) for 5-10 min at 37°C. The single-cell suspension is then seeded into stirred bioreactors (Reprocell) or spinner flasks at a density of 500k to 1 million cells per mL and spun at a rate of 65-80 rpm for 48h, leading to the spheroid formation. After forming spheroids, kidney organoids are differentiated using a directed differentiation method as previously described (Morizane and Bonventre, 2017), with adjustments in the vessel selection for stirred bioreactors or spinner flasks.

### Kidney tissue assembly

Each kidney tissue disc is assembled by depositing a densely cellular matrix composed of kidney organoid building blocks in a fibrinogen solution into cylindrical cavities stamped in an extracellular matrix composed of 8 wt% gelatin, 2.5 mM CaCl_2_, and 10 U thrombin, in DMEM/F12 medium. The gelatin is made by first preparing a 15% (w/v) gelatin solution (Type A, 300 bloom form porcine skin, Sigma) by adding prewarmed PBS without Ca^2+^ and Mg^2+^ to the gelatin powder. This gelatin solution is stirred for 12h at 70°C to allow for the complete dissolving of the gelatin. A 250 mM CaCl_2_ stock solution is made by dissolving CaCl_2_ pellets in sterile water and storing them at 4°C for long-term storage. 2,000 U/mL stock solutions of thrombin are created by reconstituting lyophilized thrombin (Sigma Aldrich) in sterile water and storing it at − 20°C. After warming all components to 37°C, the constituents are mixed together in the following order: DMEM/F12, gelatin, CaCl_2_, and thrombin. The gelatin-thrombin solution is held at 37°C for 5-10 min to ensure proper mixing prior to casting into a PDMS reservoir. Next, a disc stamp produced by stereolithography, a 3D printing method, using htm140v2 resin or Biomed clear resin and treated with 5 wt% Pluronic F127 (BASF) to prevent adhesion, is placed on top of the PDMS reservoir containing the gelatin-thrombin mixture. The stamp and PDMS reservoir containing the gelatin-thrombin solution are allowed to cool at 4°C for at least 30 min to solidify this gel. The 3D-printed stamp is then carefully removed, leaving behind the cylindrical cavities that serve as individual molds for each tissue disc. Each mold is transferred to an ultra-low adhesion 6-well plate using an 8 mm biopsy punch and spatula when they are held at 4°C for no longer than 24h prior to tissue disc casting.

Before casting, kidney organoids are collected from the stirred bioreactors using a 10mL serological pipette, passed through a 500 µm cell strainer to remove larger aggregates, and allowed to settle at the bottom of a conical tube. A fibrinogen mixture is created containing: 10mg/mL fibrinogen and 2.5 mM CaCl_2_ in DMEM/F12 media. The fibrinogen stock solution is made by first preparing a 50 mg/mL stock from lyophilized bovine blood plasma protein (Millipore). It is reconstituted in a controlled manner by adding sterile PBS without Ca^2+^ and Mg^2+^ and keeping it at 37°C without agitation for 2–3h. Once complete, the fibrinogen solution is stored in smaller aliquots at − 20°C for later use. The CaCl_2_ stock solution is described using the previously described process. Once the fibrinogen and CaCl_2_ are properly mixed into the DMEM/F12 to create the fibrinogen mixture, excess media is removed. The organoids are then mixed with 3x the volume of the fibrinogen mixture. This organoid-laden fibrinogen solution is allowed to settle for ∼2 min and the supernatant is removed. The remaining organoids are then mixed with 1x volume of fibrinogen mixture and centrifuged at 30 rcf for 3 min. 200 µL Eppendorf Combitip advanced positive displacement tips (Eppendorf) are cut with a razor blade to make a wide bore opening on the depositing end and loaded with small volumes of 5 wt% gelatin in PBS to protect the organoids from the displacement plunger. The tips are loaded with the organoid-fibrinogen and then centrifuged in a custom 3D printed holder at 30 rcf for 3 min to create a compacted organoid-fibrinogen tissue matrix. Once compacted, an Eppendorf Repeater M4 multi-dispenser pipette is used to deposit 8 µL of this compacted tissue matrix into each disc mold, where it resides for ∼5 min to allow thrombin diffusion from the surrounding gelatin matrix that drives polymerization of the fibrin gel, i.e., which “binds” the organoids together. A droplet of the gelatin-thrombin mixture is then placed on top of each disc to ensure the polymerization of that region. 5 mL of Advanced RPMI media with 1% GlutaMAX, 1.5% FBS, 1% anti-anti, and 1% aminocaproic acid (ACA) is then added to each well and placed on an orbital shaker at 90 rpm in the incubator. Full-volume media changes are performed 24h after disc formation and each subsequent 48h over a 7-day culture period prior to implantation. The kidney tissue discs are live-imaged at days 0, 4, and 7 using a Keyence zoom microscope (VHX-2000, Keyence, Japan).

### Mice

NSG (NOD.Cg-Prkdc^scid^ Il2rg^tm1Wjl^/SzJ) mice were purchased from the Jackson Laboratory and maintained as breeding colonies in the Massachusetts General Hospital animal facility with water and food ad libitum. All mice used in the experiments were 8 to 12 weeks old. Following the Institutional Animal Care and Use Committee (IACUC) and National Institutes of Health (NIH) Animal Care guidelines, mice were bred and housed in individual and standard mini-isolators under specific pathogen-free conditions. The Mass General Brigham IACUC approved all experiments under protocol number 2020N000125.

### Peripheral blood mononuclear cells (PBMCs) isolation

PBMCs were used as allogeneic human immune cells. PBMCs were isolated from peripheral blood samples of a healthy volunteer by density gradient centrifugation using Ficoll-Paque solution (GE Healthcare) and were freshly transferred to NSG mice. These experiments were approved under the Mass General Brigham IRB number 2017P000298.

### HLA typing and eplet mismatches calculations

Allogeneic human immune cells (PBMCs) and iPSCs (BJFF cells) were HLA typed by the Histogenetics company using a 4x resolution (class I: exons 1-8 typed, and class II exons 2-6 typed). Eplets mismatches were calculated using the calculator from the HLA Eplet Registry (https://www.epregistry.com.br/calculator/index.html).

### Kidney tissue implantation

NSG recipient mice were subcutaneously transplanted with ∼10 mm^3^ human kidney organoid discs. Discs were surgically positioned into the fat pad on the abdomen. On the next day, 13 x 10^6^ freshly isolated human PBMCs (allogeneic human immune cells) were transferred to tissue-implanted NSG mice via retro-orbital injection of the venous sinus. One cohort of NSG mice was transplanted with the kidney organoid discs but did not receive allogeneic human immune cells (PBMCs) as a negative control. Transplanted kidney tissues and spleens were procured on days 4, 14, 21 and 28 after the transfer of allogeneic human immune cells (**Suppl. Fig 1**) for flow cytometry, histology and transcriptional analyses.

### Flow cytometry

We stained the isolated kidney cells and splenocytes for flow cytometry. Isolated cells were mouse (TruStain FcX, anti-mouse CD16/32, clone 93, Biolegend) and human (TruStain FcX, Fc receptor blocking solution, Biolegend) Fc-blocked for 20 min before staining for surface markers for 30 minutes in FACS buffer (2% FBS in PBS 1x) on ice. We used the following anti-human antibodies: CD8-BUV737 (1:50, clone SK1, BD Biosciences), CD4-BUV395 (1:50, clone SK3, BD Biosciences), CD19-BV786 (1:50, clone HIB19, Biolegend), CD123-BV711 (1:20, clone 6H6, Biolegend), CD27-BV650 (1:50, clone O323, Biolegend), CD3-BV605 (1:50, clone OKT3, Biolegend), CD14-BV510 (1:25, clone M5E2, Biolegend), CD66b-BV421 (1:50, clone 6/40c, Biolegend), CD127-PerCP-Cy5.5 (1:20, clone A019D5, Biolegend), CD16-FITC (1:40, clone 368, Biolegend), CD56-PE-Cy7 (1:50, clone 5.1H11, Biolegend), CD45-PE-Cy5 (1:100, clone HI30, Biolegend), CCR7-PE-CF594 (1:20, clone G043H7, Biolegend), CD25-PE (1:20, clone M-A251, Biolegend), HLA-DR-APC-Cy7 (1:50, clone L243, Biolegend), HLA-DR-BV650 (1:50, clone L243, Biolegend), CD11c-AF700 (1:25, clone 3.9, Biolegend), CD45RA-APC (1:50, clone HI100, Biolegend), HLA-A/B/C-BV605 (1:20, clone W6/32, Biolegend). We used the following anti-mouse antibodies: CD45-BV421 (1:100, clone 30-F11, Biolegend). After surface staining, cells were stained for Annexin V-BV510 (1:50, Biolegend) in Annexin V binding buffer (Biolegend) for 15 min at room temperature. Stained cells were analyzed on a LSR Fortessa X-20 flow cytometer (BD Biosciences) with FACSDiva software (BD Biosciences). Data were analyzed with FlowJo software (TreeStar). Viable cells were selected based on the staining with LIVE/DEAD Fixable Blue Dead Cell Stain Kit (Thermo Fisher) prior to the Fc-blocking.

### Immunohistochemistry and immunofluorescence imaging

All specimens were formalin-fixed, processed and paraffin-embedded. Five μm-thick sections were prepared and stained with hematoxylin and eosin or were used for immunohistochemical studies. For some stains (i.e., CD31), immunohistochemical staining was performed using an automated Ventana BenchMark Stain System (Roche). Antibodies used were rabbit polyclonal anti-human CD31 (Abcam). Stained sections were photographed using an Axio Imager M2 microscope (Zeiss) and processed with ImageJ and Photoshop software.

Other antibodies used were rabbit monoclonal anti-human podocalyxin (anti-PODXL, Abcam, clone EPR9518), rabbit monoclonal anti-e-cadherin (anti-CDH1, Cell Signaling, clone 24E10), mouse monoclonal anti-human CD4 (Biocare, clone 4B12), mouse monoclonal anti-human CD8 (Biocare, clone C8/144B), mouse monoclonal anti-human CD31 (Agilent, clone DAKO JC70A), mouse monoclonal anti-HLA class II-DR/DP/DQ (Abcam, clone CR3/43), mouse monoclonal anti-HLA class I A/B/C (Abcam, clone EMR8-5). Photomicrographs were taken using an Olympus BX53 microscope with a DP27 camera and cellSens Imaging Software (Olympus Life Science Solutions).

We used a Leica Zeiss LSM 880 + FLIM microscope to carry out all immunofluorescence confocal imaging. Harvested tissue discs were formalin-fixed and placed overnight at 4°C in a blocking solution containing 0.125% Triton X-100 and 1 wt% donkey serum in DPBS with calcium and magnesium. The blocking solution was washed off using DPBS, and primary antibodies were added for 24-48 hours at 4°C in a staining solution containing 0.5 wt% BSA and 0.125% Triton X-100 in DPBS. Primary antibodies were washed off using DPBS and secondary antibodies were added using the staining solution described above. The primary antibody list used for the stains is provided in **Suppl. Table 2**. DAPI (405), Alexa Fluor 488, Alexa Fluor 555, or Alexa Fluor 647 were used for secondary antibodies. Fibrin confocal imaging was conducted by imaging autofluorescence in the 555 channel, which was coupled with the PODXL antibody. Image J software was used for confocal image processing.

### RNA extraction

We obtained six consecutive 10 μm sections from formalin-fixed paraffin-embedded (FFPE) kidney tissues harvested at different time points. Deparaffinization with Xylene and RNA extraction was performed in sterile 1.5 ml microcentrifuge tubes with the RNeasy FFPE Kit (Qiagen), according to the manufacturer’s instructions. The concentration and purity of the isolated total RNA were measured using the 2100 Bioanalyzer system (Agilent) at the Center for Advanced Molecular Diagnostics (CAMD) Research Core of the Brigham and Women’s Hospital. The absorbance ratio at 260/280 was used to determine RNA quality.

### NanoString nCounter assay for mRNA gene expression assay

We analyzed 770 genes with the NanoString nCounter PanCancer Immune Profiling Gene Expression (GX) Codeset. Gene expression was measured on 100–200 ng of extracted RNA. Samples were processed on the NanoString nCounter Analysis System (NanoString Technologies) following the manufacturer’s instructions at the CAMD Research Core of the Brigham and Women’s Hospital. We normalized raw gene expression counts and performed background correction, quality control and analyses with the nSolver Analysis Software (Version 4.0.70). Ten reference genes (*POLR2A*, *SDHA*, *SF3A1*, *TMUB2*, *NRDE2*, *MRPL19*, *PUM1*, *DNAJC14*, *STK11IP* and *ERCC3*) were used for normalization. We used the quality control parameters recommended by the manufacturer.

To compare our gene signature with transcripts in acute cellular rejection in human kidney graft biopsies, we selected the upregulated genes from Win *et al*. (Win et al., 2021) and van der Zwan *et al*. (Van Der Zwan et al., 2019) and intersected them with the upregulated DEGs from this study. We defined upregulated genes as those with log2 fold change greater than 1.5 and a p-value less than or equal to 0.05. Some analyses were performed with Python 3.9.16, pandas 1.3.5 and matplotlib 3.7.1.

### Statistical Analyses

For flow cytometry, t-SNE and FlowSOM analyses were performed on FlowJo (v.10.8.1). Cell populations were downsampled from viable singlets human CD45^+^ cells using the Downsample (v.3.3) plugin for Flowjo, followed by concatenation. We then performed FlowSOM clustering on the concatenated file with the FlowSOM plugin (v2.6) was performed on the concatenated file with n = 8-9 clusters for the human immune cell populations. The FlowSOM output was surface markers expression and the percentage of cells in each cluster was used to calculate absolute numbers. The t-SNE maps were built with a vantage point tree algorithm with 1000 iterations. Cluster identification in the map was performed with the help of the Cluster Explorer plugin (v1.6.3) through the heatmap expression of surface markers CD16, CD127, CD66b, CD14, CD3, CD27, CD123, CD19, CD4, CD8, CD45RA, CD11c, HLA-DR, CD25, CCR7, and CD56.

## Acknowledgments

This work was supported by the Wellcome LEAP Human Organ Physiological Engineering (HOPE) program, the NIH Re(Building) a Kidney Consortium (NIH UC2DK126023), the NIH award DP2EB029388/DK133821 to R.M., NIH grants R56DK122380 and UC2DK126023 to R.M., and R01AI143887-05 to L.V.R. We thank P. Stankey for assistance in printing the 3D stamps. A. Moisan, K. Wolf, K. Kroll, D. Reynolds, N. Gupta and M. Creighton for insightful discussions and the Harvard Center for Biological Imaging for experimental assistance, and Kazumi Ida and Chengcheng Zhang for technical assistance in kidney organoid differentiation.

## Contributions

T.J.B., J.O.A., R.V.G., S.U., R.M., J.A.L, and L.V.R. designed the research. T.J.B., Y.G., and R.B.G. performed disc implantation, harvesting, tissue processing, and flow cytometry experiments. T.J.B. and K.L. analyzed the flow cytometry experiments. T.J.B. and G.T.B. performed transcriptomics experiments and analyses. T.J.B., J.O.A., K.K., M.T., K.H. and I.R. performed histological analyses. J.O.A., K.K., M.T., K.H. and R.M. developed the direct 3D differentiation approaches using stirred bioreactors. J.O.A., K.K., M.T., K.H. and J.E.R. generated kidney organoids for the discs. J.O.A. and S.U. conceived, designed, and implemented the tissue disc stamping approach used to create uniform discs. J.O.A. and R.V.G. casted, cultured, and characterized the tissue discs prior to implantation. T.J.B., Y.G., J.O.A., J.A.L. and L.V.R. wrote the manuscript. All authors edited the manuscript.

