## Supplemental Figures and Tables for "Immune Response of Transplanted Kidney Tissues Assembled from Organoid Building Blocks"

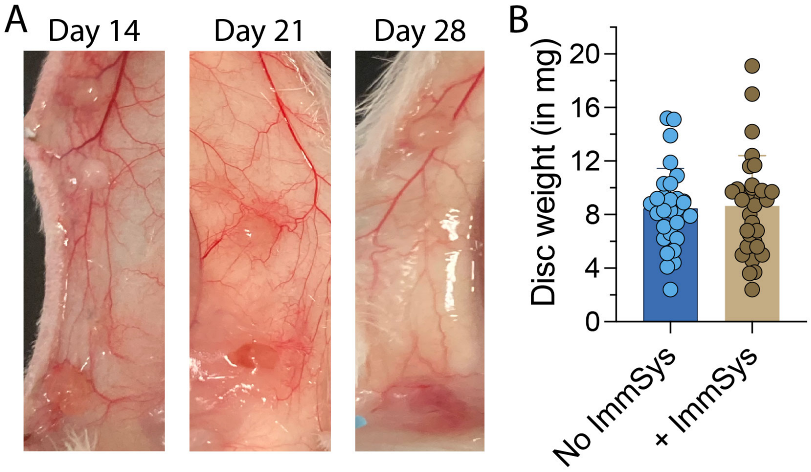


**Supplemental Figure 1. Kidney tissues harvested after transplantation.** (**A**) Images of the kidney tissues that were subcutaneously implanted into the NSG mice reconstituted with allogeneic human immune cells, and harvested on days 14, 21, and 28. (**B**) Weights (in mg) of kidney tissues harvested from NSG mice reconstituted with (+ ImmSys) or without (No ImmSys) allogeneic human immune cells. Data are representative of two pooled experiments with all the time points in the same experimental group (n = 30 - 32 tissues/group). Statistics are performed using an unpaired t-test.

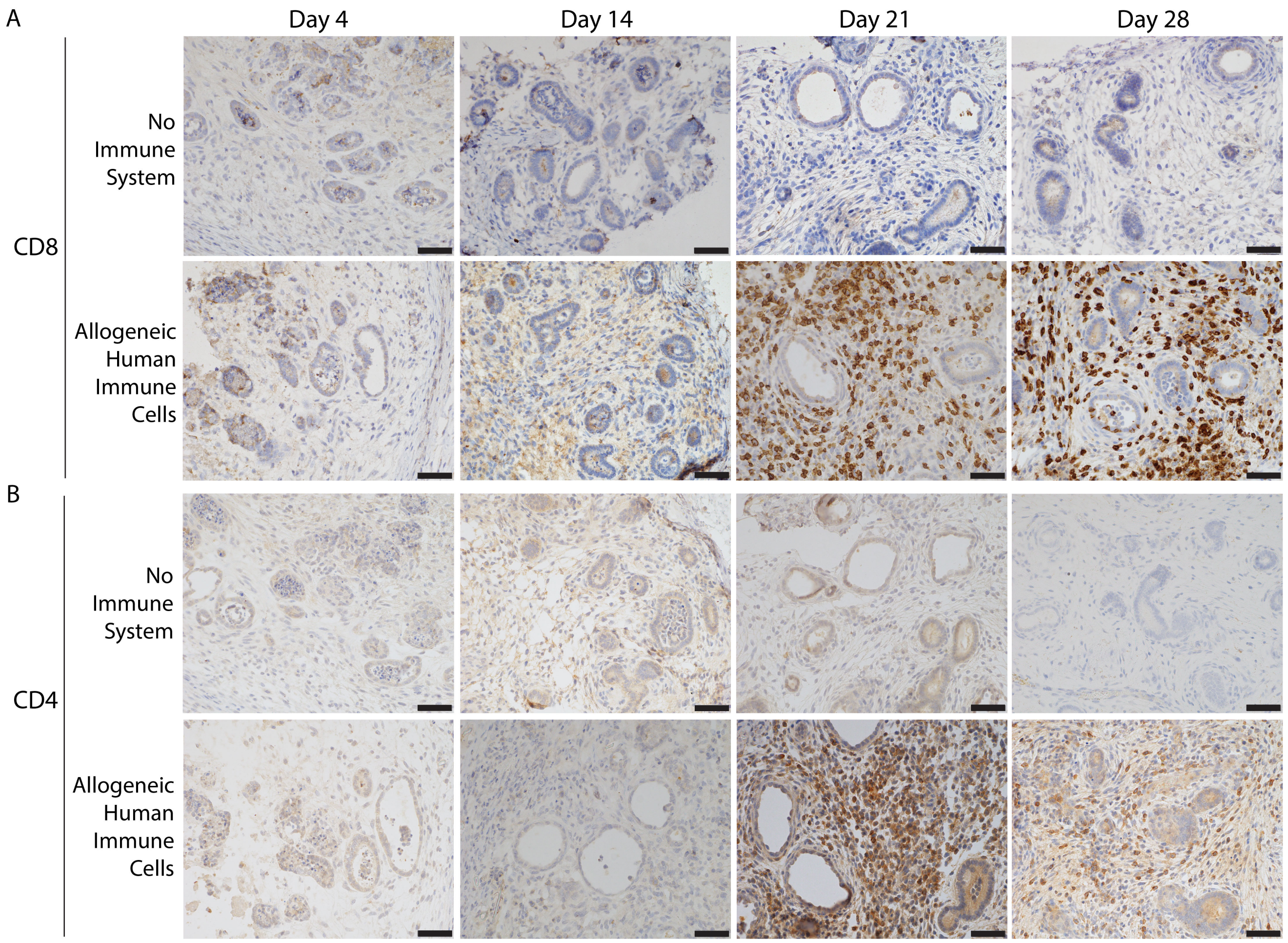


**Supplemental Figure 2. Human kidney disc infiltration by CD8⁺ and CD4⁺ cells.** Kidney tissues are transplanted into NSG mice reconstituted with allogeneic human immune cells, harvested and then analyzed on days 4, 14, 21 and 28. (**a**) Immunostaining of (**A**) CD8, and (**B**) CD4 over time. Data are representative of two independent experiments. Scale bars = 50 μm.

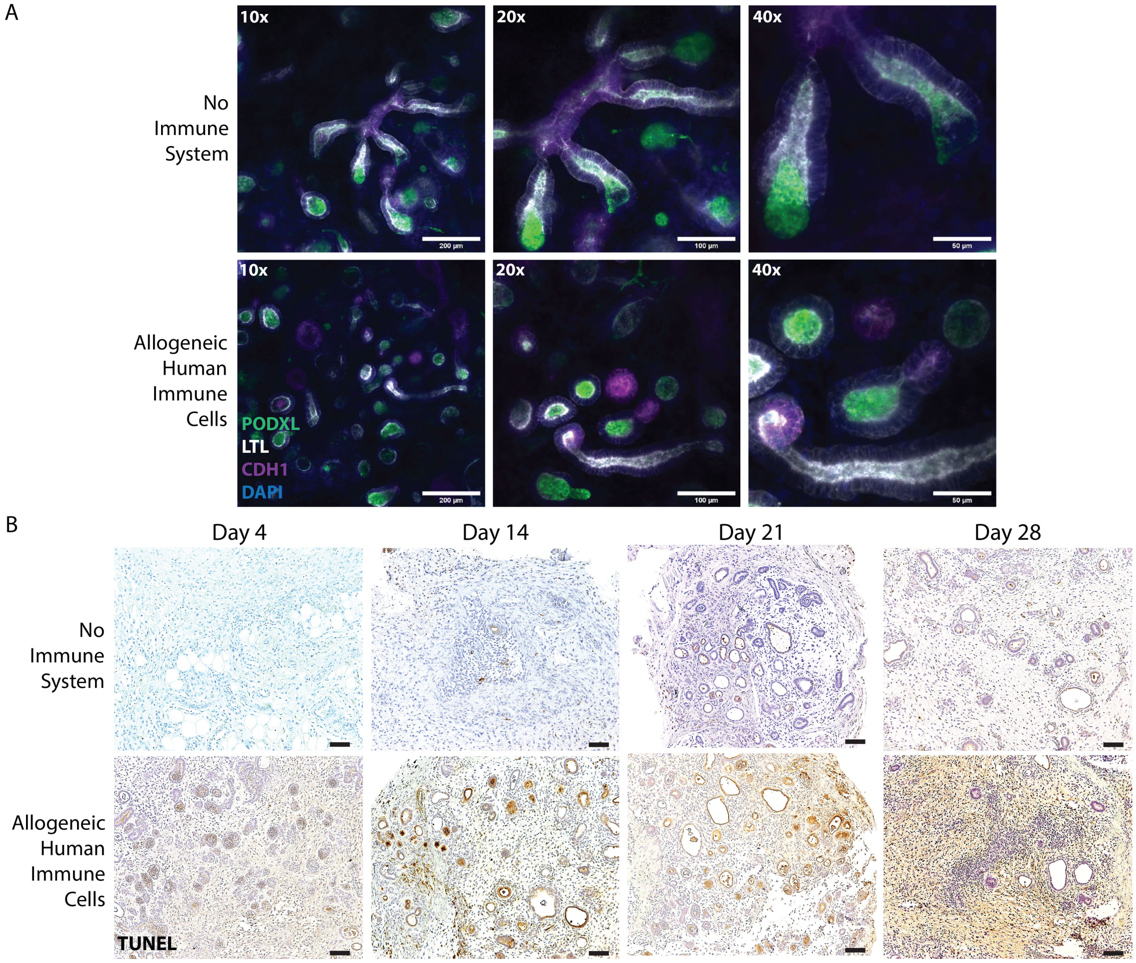


**Supplemental Figure 3. Allogeneic human immune cells induce kidney tissue apoptosis *in vivo*.** Kidney tissues are transplanted into NSG mice reconstituted with allogeneic human immune cells, harvested and then analyzed on days 4, 14, 21 and 28. (**A**) Confocal images of PODXL (glomeruli, green), LTL (proximal tubules, gray), CDH1 (distal tubules, purple) and DAPI (nuclei, blue) expression by tissues harvested on day 21. Scale bars = 1 mm (tiled disc overview), 200 µm (10x), 100 µm (20x), and 50 µm (40x). Data are representative of two independent experiments. (**B**) Kidney tissue TUNEL immunostaining over time. Data are representative of three independent experiments. Scale bars = 100 µm.

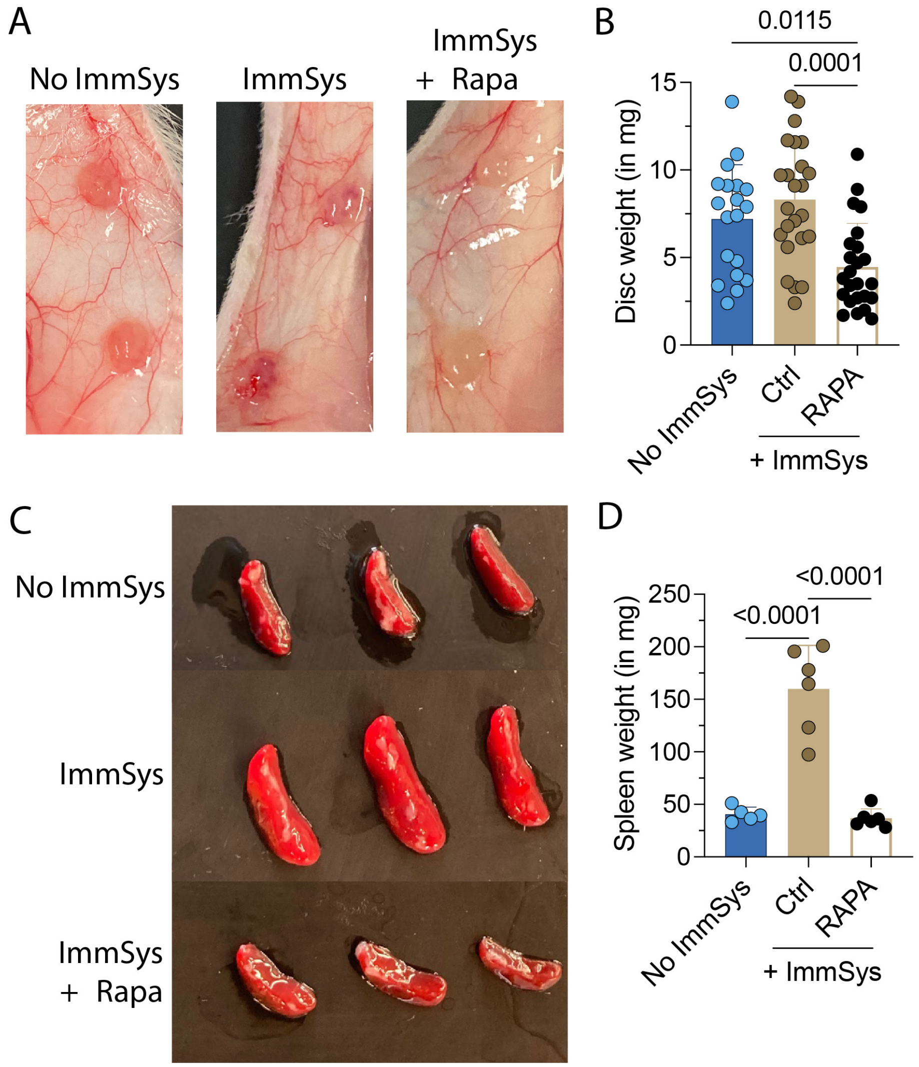


**Supplemental Figure 4. Kidney tissues and spleens harvested after transplantation.** Kidney tissues were transplanted into NSG mice reconstituted with (ImmSyst) or without (No ImmSyst) allogeneic human immune cells. One cohort of NSG mice was treated with 200 mg/kg/day of Rapamycin (ImmSyst + Rapa). (**A**) Images and (**B**) weights (in mg) of kidney transplanted tissues harvested on day 21 after the transfer of allogeneic human immune cells. Data are representative of two pooled experiments. Statistic by one-way ANOVA with Tukey post-test (n = 19 - 24 tissues/group). (**C**) Visual aspects and (**d**) weights (in mg) of spleens on day 21 after the transfer of allogeneic human immune cells. Data are representative of two pooled experiments. Statistic by one-way ANOVA with Tukey post-test (n = 6 spleens/group).

### Supplemental Tables

#### Suppl. Table 1. HLA typing of iPSCs and immune cells

| | iPSCs | Human Immune Cells |
|---|---|---|
| *HLA-A* | *A*01:01*<br>*A*01:01* | *A*25:01*<br>*A*42:02* |
| *HLA-B* | *B*08:01*<br>*B*41:01* | *B*18:01*<br>*B*44:03* |
| *HLA-DRB1* | *DRB1*03:01*<br>*DRB1*07:01* | *DRB1*07:01*<br>*DRB1*15:01* |

#### Suppl. Table 2. Primary antibodies used for immunofluorescence confocal imaging

| **Target** | **Clone** | **Concentration** | **Secondary** |
|---|---|---|---|
| PODXL | AF1658 | 1:300 | Anti-Goat |
| LTL | B-1325 | 10mg/mL | Biotinylated |
| CDH1 | ab40772 | 1:300 | Anti-Rabbit |
| PECAM1 (CD31) | ab9498 | 1:300 | Anti-Mouse |
